# Suppressing Primer-Driven Nonspecific Amplification in LAMP Using TrueLAMP

**DOI:** 10.1101/2025.10.23.684253

**Authors:** Zhidong Chen, Hyuck-Jin Seo, Paul Collier

**Affiliations:** Boltii Diagnostics Inc., Salt Lake City, Utah, USA; Department of Molecular Biosciences, University of Texas at Austin, Austin, Texas, USA; Department of Physiology and Biophysics, Weill Cornell Medicine, New York, New York, USA

**Keywords:** Colorimetric LAMP, Loop-mediated isothermal amplification, no-template control, nonspecific primer amplification, false positive, point-of-care diagnostics, TrueLAMP™

## Abstract

Loop-mediated isothermal amplification (LAMP) enables rapid nucleic acid detection but is vulnerable to primer-driven nonspecific amplification that can produce false-positive results. We developed TrueLAMP, a colorimetric LAMP formulation containing an inhibitor that suppresses nonspecific amplification while preserving target-dependent amplification. All 65 no-template controls across these studies remained negative, corresponding to 100% observed specificity. Using SARS-CoV-2 as a model and the Lamb primer set as a representative benchmark, TrueLAMP achieved a limit of detection of 250 copies per reaction with 95% detection probability. Endpoint color remained stable during extended incubation, reducing the need for time-critical readout. TrueLAMP was compatible with readily available heating devices, including a convection oven, water bath, and temperature-controlled smart mug. Independent evaluations in two external laboratories using additional primer–template systems, including DNA targets, further supported suppression of nonspecific amplification and sequence-dependent target detection. Together, these findings support TrueLAMP as a practical approach for improving the reliability of LAMP detection while retaining a simple isothermal workflow.

**Method summary:** Colorimetric LAMP reactions targeting SARS-CoV-2 were performed using TrueLAMP polymerase and buffer containing a nonspecific amplification inhibitor. Reactions used a two-step incubation consisting of 60 °C for 10 min followed by 65 °C and were heated in a convection oven, water bath, or temperature-controlled smart mug. Amplification was assessed by visual color change and quantified using smartphone-based magenta or green channel measurements. Agarose gel electrophoresis provided orthogonal confirmation of amplification products in reactions with and without the inhibitor. Independent external-laboratory evaluations used additional primer–template systems to assess suppression of nonspecific amplification and sequence-dependent target detection.

**ARTICLE HIGHLIGHTS:**

- Across multiple primer sets and experimental conditions, all 65 TrueLAMP no-template controls remained negative (100% observed specificity, exact 95% CI, 94.5–100%).
- Using the Lamb primer set, TrueLAMP achieved a limit of detection of 250 RNA copies per reaction with 95% detection probability (95% CI, 77–99%).
- Agarose gel analysis confirmed suppression of nonspecific amplification products while preserving target-dependent amplification.
- Stable endpoint detection and a two-step 60 °C → 65 °C protocol supported reliable operation across readily available heating devices.
- Independent evaluations in two external laboratories supported suppression of nonspecific amplification and sequence-dependent target detection.

## 1. Introduction

Loop-mediated isothermal amplification (LAMP) is widely used for rapid nucleic acid detection because it provides efficient amplification under isothermal conditions and can be implemented with relatively simple equipment [1–4]. These features make LAMP attractive for point-of-care and field testing. A persistent limitation, however, is nonspecific amplification in the absence of target nucleic acid. Such amplification can produce false-positive signals in no-template controls (NTCs) and potentially in test samples, reducing assay reliability [5–7].

False-positive LAMP reactions are often attributed first to contamination. Investigators may replace reagents and consumables, separate preparation and amplification areas, or introduce uracil-DNA glycosylase (UDG) and dUTP systems to reduce carryover contamination. If false positives persist, primer sets are often redesigned and rescreened. These steps can add substantial time and complexity to assay development when the underlying problem is primer-driven amplification rather than contamination.

Additives including betaine, DMSO, tetramethylammonium chloride (TMAC), and pullulan have been evaluated as ways to improve LAMP specificity [5, 7–9]. These approaches can reduce nonspecific amplification under some conditions, but suppression may depend on primer set, formulation, and incubation time. Delayed nonspecific amplification remains particularly problematic when reactions are incubated long enough to improve detection of low-copy targets.

Other approaches couple LAMP to sequence-specific nucleases such as CRISPR/Cas12a or programmable Argonaute (pAgo) proteins. These systems use sequence recognition after or alongside amplification to distinguish target-derived products from off-target or primer-derived products [10–13]. They can improve analytical specificity, but they also add reagents, guide design, and detection steps to the assay.

We hypothesized that a substantial fraction of nonspecific LAMP amplification arises from primer– primer interactions, self-priming structures, and other mispriming events that can be extended by strand-displacing polymerases. Mechanistic studies have shown that false-positive NTC signals can arise from primer dimers, self-amplifying hairpins, and related primer-derived products [14–15].

We developed TrueLAMP, a polymerase formulation containing a proprietary inhibitor designed to suppress primer-driven nonspecific amplification while retaining target-dependent amplification. The inhibitor acts within the amplification reaction rather than through a separate sequence-specific detection layer. This approach is intended to preserve the operational simplicity of LAMP while extending the interval over which negative reactions remain stable.

Here, we evaluated TrueLAMP using multiple independently designed LAMP primer sets, SARS-CoV-2 RNA and heat-inactivated virions, colorimetric time-course analysis, and agarose gel electrophoresis. We tested whether primer-driven nonspecific amplification could be suppressed while retaining target detection and analytical sensitivity. Key specificity findings were also evaluated independently in two external laboratories using additional primer–template systems.

## 2. Materials and methods

### 2.1. Reagents and Enzymes

TrueLAMP™ kits (catalog # TL1001M) were obtained from Boltii Diagnostics Inc. (Salt Lake City, UT, USA), comprising TrueLAMP polymerase and 2x TrueLAMP buffer containing an inhibitor that suppresses nonspecific LAMP amplification. The inhibitor composition is proprietary and is not disclosed. SARS-CoV-2 RNA (NR-52347) and heat-inactivated SARS-CoV-2 virus (NR-52286) were obtained from BEI Resources, NIAID, NIH.

For evaluation in the Ellington Laboratory, complete genomic DNA isolated from *Salmonella enterica* subsp. *enterica* serovar Typhi strain Ty2 (GenBank accession AE014613.1, NR-543) was obtained from BEI Resources, NIAID, NIH.

For evaluation in the Mason Laboratory, inactive gBlock DNA fragments representing Shamonda virus (SHV, N and Ns genes, NC_018464.1) and avian myelocytomatosis virus (AMV, NC_001866.1) sequences were obtained from Integrated DNA Technologies (IDT).

### 2.2. Primer Sets

Four independent LAMP primer sets targeting SARS-CoV-2 genomic RNA were used for the primary characterization of TrueLAMP: one proprietary set (N008) from Boltii Diagnostics and three published sets—Lamb, N27, and M3. The Lamb primer set corresponds to the SARS-CoV-2 RT-LAMP design reported by Lamb et al. (2020) [16], whereas N27 and M3 were adopted from the LAMP optimization study of Huang, Tang, Ismail, and Wang (2022) [17].

Two additional published primer sets, HJS-18 and HJS-13-6 [18], were used for evaluation in the Ellington Laboratory. HJS-18 was included as a challenge primer set because of its previously reported propensity for rapid nonspecific amplification in NTC reactions. HJS-13-6, targeting the STY2879 gene of *S*. Typhi, was used to evaluate target-dependent amplification and suppression of NTC amplification.

Primer sets used in the Mason Laboratory were designed in-house against loci represented in the SHV and AMV gBlock controls. Primers were synthesized by IDT Custom DNA Oligo service.

Full primer sequences are provided in Supplementary Table S1. Published sequences used in this study matched those reported in the corresponding original publications. Unless otherwise specified, final primer concentrations in the 10 µL reaction were 0.2 µM F3/B3, 0.4 µM LF/LB, and 1.6 µM FIP/BIP.

### 2.3. LAMP Reactions

Reactions were performed as described below, with additional procedural details provided in Supplementary Protocol S1. Each nominal 10 µL reaction contained 3.0 µL nuclease-free water, 5.0 µL 2x TrueLAMP buffer, 0.05 µL TrueLAMP polymerase, 1.0 µL 10x primer mix, and either 1.0 µL nuclease-free water for the no-template control (NTC) or 1.0 µL template (RNA, DNA, or heat-inactivated virions). Master mixes were prepared with approximately 10% excess volume to compensate for pipetting and dispensing losses.

Unless otherwise indicated, reactions were incubated using a two-step protocol consisting of 60 °C for 10 min followed by 65 °C for 80 min. Positive reactions changed from red to orange/yellow, whereas negative reactions remained red.

Colorimetric signals were quantified by measuring either maximum magenta intensity in CMYK mode or minimum green intensity in RGB mode using a smartphone colorimeter application. Measurements were taken immediately below the liquid meniscus using the smallest available sampling aperture. Both metrics reflect the same underlying red-to-yellow color transition during amplification.

### 2.4. Colorimetric LAMP Setups

Because the reactions were performed in a nominal 10 µL volume, uniform heating around the reaction tubes was important to limit evaporation and condensation. Representative heating and imaging configurations are shown in Figure 1. For semi-real-time LAMP, 8-strip PCR tubes were placed upright in a convection hybridization oven (Stratagene PersonalHyb, Model 401030) and photographed at 15-min intervals through the viewing window with a fixed-position smartphone. Imaging geometry and illumination were kept constant because changes in angle, distance, or lighting affect numerical color measurements. For endpoint LAMP, reactions were heated in a laboratory water bath or a temperature-controlled smart mug. Temperature at the tube position was verified with a liquid thermometer or thermocouple.

**Figure 1.**
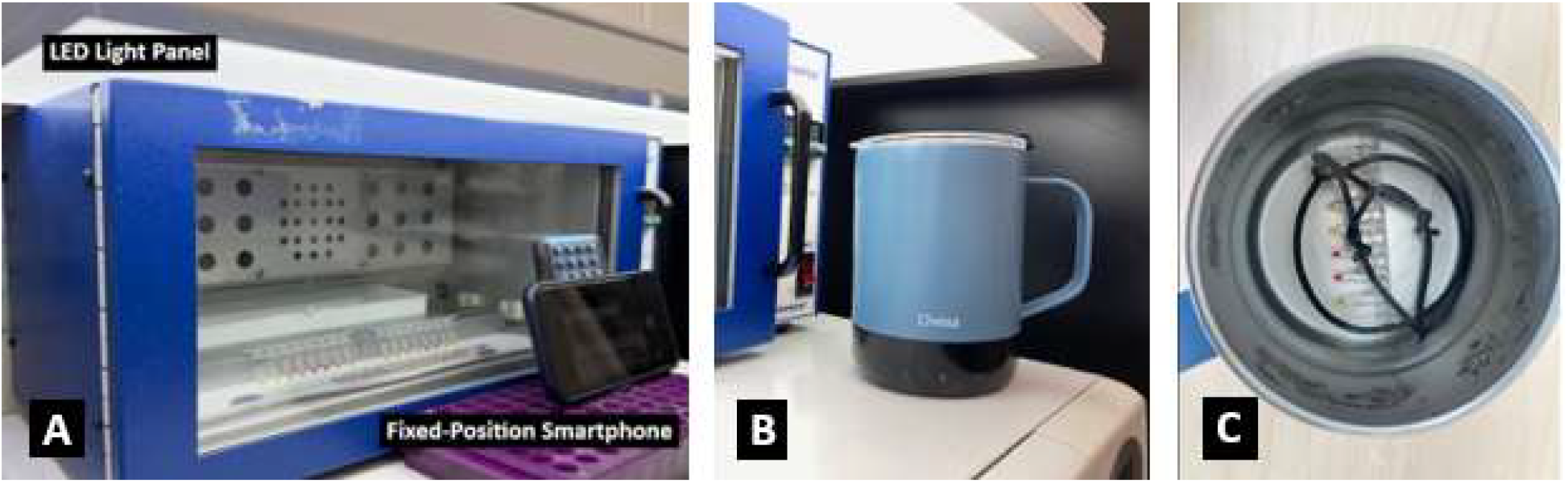
Semi-real-time and endpoint colorimetric LAMP setups using off-the-shelf equipment. (A) Semi-real-time monitoring in a convection oven using fixed-position smartphone imaging under constant illumination. (B) Temperature-controlled smart mug used as an endpoint incubator. (C) Interior view showing the 8-strip reaction tubes held down to maintain immersion and consistent thermal contact. These configurations show that TrueLAMP can be performed with unmodified, readily available equipment.

### 2.5. Independent External-Laboratory Evaluation

TrueLAMP was independently evaluated in the Ellington Laboratory at the University of Texas at Austin and the Mason Laboratory at Weill Cornell Medicine. These experiments used primer–template systems distinct from the SARS-CoV-2 assays and examined complementary aspects of reaction specificity.

#### 2.5.1. Ellington Laboratory evaluation

Two previously characterized *S*. Typhi LAMP primer sets, HJS-18 and HJS-13-6 [18], were evaluated in the Ellington Laboratory. HJS-18 was included specifically as a challenge for primer-driven nonspecific amplification because it had previously shown rapid NTC amplification. HJS-13-6 was used to assess whether NTC suppression could be maintained while retaining target-dependent amplification.

Complete genomic DNA from *S*. Typhi strain Ty2 (NR-543, BEI Resources, NIAID, NIH) was used as template. The DNA was resuspended in TE buffer to 1 × 10^7^ genome copies/µL, aliquoted, and stored at −20 °C. For positive reactions, DNA was diluted to 2,000 genome copies/µL, and 1 µL was added to each nominal 10 µL reaction, corresponding to 2,000 genome copies per reaction. NTC reactions received 1 µL nuclease-free water. HJS-18 and HJS-13-6 reactions were run in parallel for 120 min, at 60 °C for the first 10 min followed by 65 °C for the remaining 110 min.

#### 2.5.2. Mason Laboratory evaluation

TrueLAMP was independently evaluated in the Mason Laboratory using matched and mismatched primer–template combinations to assess sequence-dependent amplification.

Inactive AMV and SHV gBlock DNA controls were used as templates. DNA concentrations were converted to copy number using the NEB BioCalculator and diluted from 10,000 to 0.1 copies/µL. Each nominal 10 µL matched reaction contained 1 µL diluted template and 1 µL of the corresponding primer mix. Mismatched reactions contained the same amount of template with a primer mix targeting the alternative gBlock sequence. NTC reactions received 1 µL nuclease-free water in place of template. AMV and SHV reactions were incubated according to the general protocol in a thermocycler with heated lid for a total 120 min.

### 2.6. Statistical analysis

Quantitative colorimetric data are presented as mean ± standard deviation. Magenta intensity between groups was compared over time using Welch’s t-test [19] and the Mann–Whitney U test [20].

Differences were considered statistically significant at p < 0.05. Fisher’s exact test [21] was used for binary outcome data. Exact 95% confidence intervals for proportions were calculated using the Clopper– Pearson method [22]. Analyses were performed in GraphPad Prism.

## 3. Results and discussion

### 3.1. Blocking Primer-Driven Nonspecific Amplification

To evaluate whether TrueLAMP suppresses primer-driven nonspecific amplification, no-template control (NTC) reactions were performed using the amplification-prone M3 primer set under identical conditions with conventional Bst polymerase (without inhibitor) or TrueLAMP polymerase (with inhibitor) at 65 °C. Magenta intensity was monitored every 15 minutes for 120 minutes (Figure 2). In inhibitor-free reactions, magenta intensity decreased progressively from 51.5 ± 1.3 at 15 min to 22.3 ± 1.5 at 120 min, indicating progressive color loss consistent with nonspecific amplification. In contrast, TrueLAMP reactions maintained stable red color throughout incubation, with magenta intensities remaining between 55.3 ± 1.0 and 54.5 ± 3.2.

**Figure 2.**
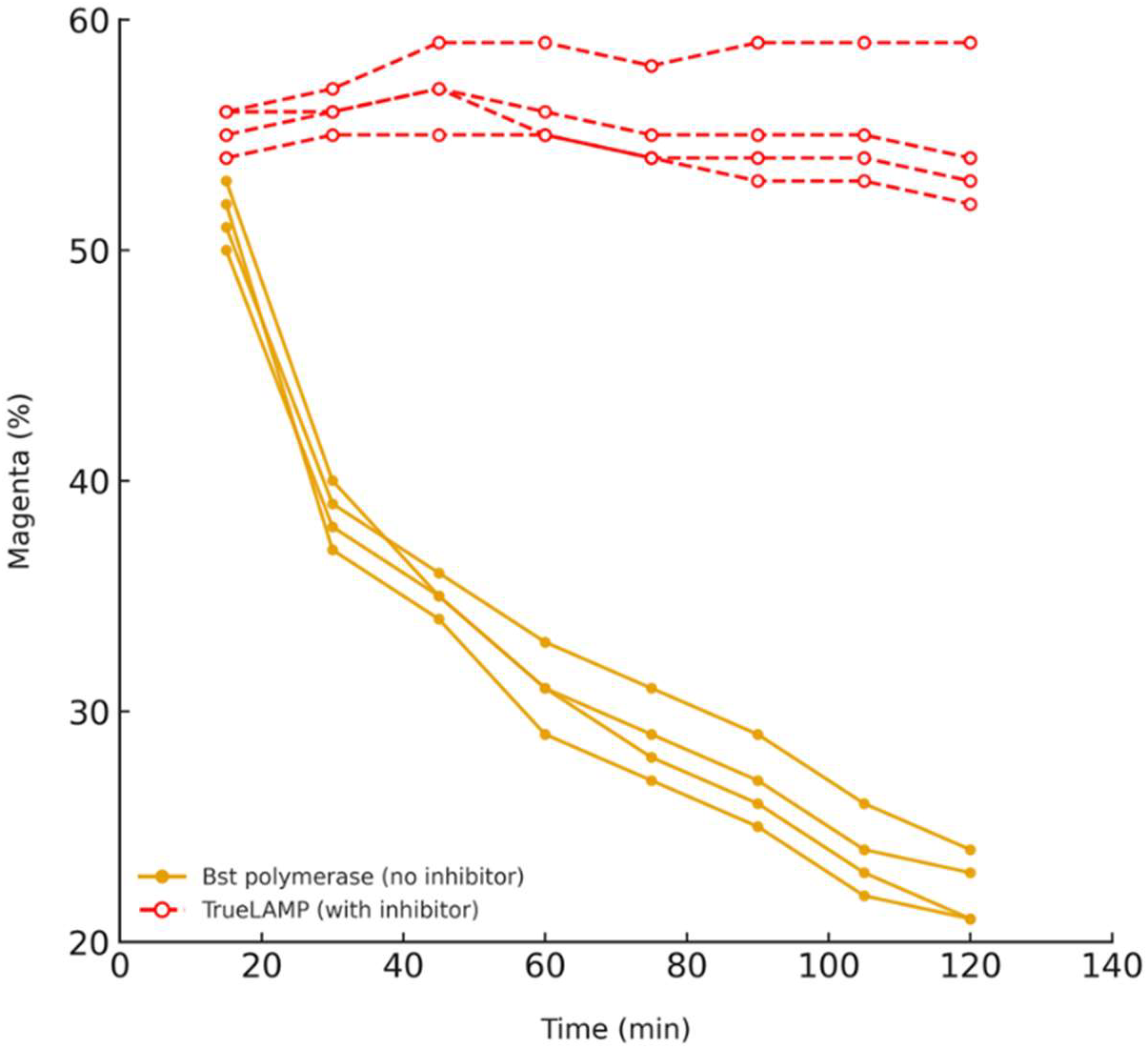
Suppression of primer-driven nonspecific amplification in no-template controls. Time-course colorimetric LAMP reactions were performed with the amplification-prone M3 primer set in the absence of template, either without inhibitor or with the TrueLAMP inhibitor. Inhibitor-free reactions showed progressive loss of magenta intensity consistent with nonspecific amplification, whereas inhibitor-containing reactions remained stable throughout the 120-min incubation.

The two conditions differed significantly at all measured time points (Welch’s t-test: p < 0.005 at 15 min and p < 0.0001 from 30 min onward). The Mann–Whitney U test gave consistent results (p ≈ 0.0286). Thus, the M3 primer set produced progressive NTC amplification without the inhibitor, whereas the TrueLAMP NTCs remained stable throughout the 120-min observation period.

Primer-driven nonspecific amplification is a recognized limitation of LAMP and can arise from primer– primer interactions and self-priming structures that are extended by strand-displacing polymerases. The stable color of the TrueLAMP NTCs shows that the formulation suppressed these events under the conditions tested.

### 3.2. Orthogonal Validation of Amplification Products

To verify that the color changes reflected nucleic acid amplification rather than indicator artifacts, reaction products were analyzed by agarose gel electrophoresis using the same polymerase and colorimetric buffer background with or without the nonspecific amplification inhibitor (Figure 3). Reactions used the M3 primer set with heat-inactivated SARS-CoV-2 virions (NR-52286) or NTCs and were incubated for 90 min in a temperature-controlled smart mug.

**Figure 3.**
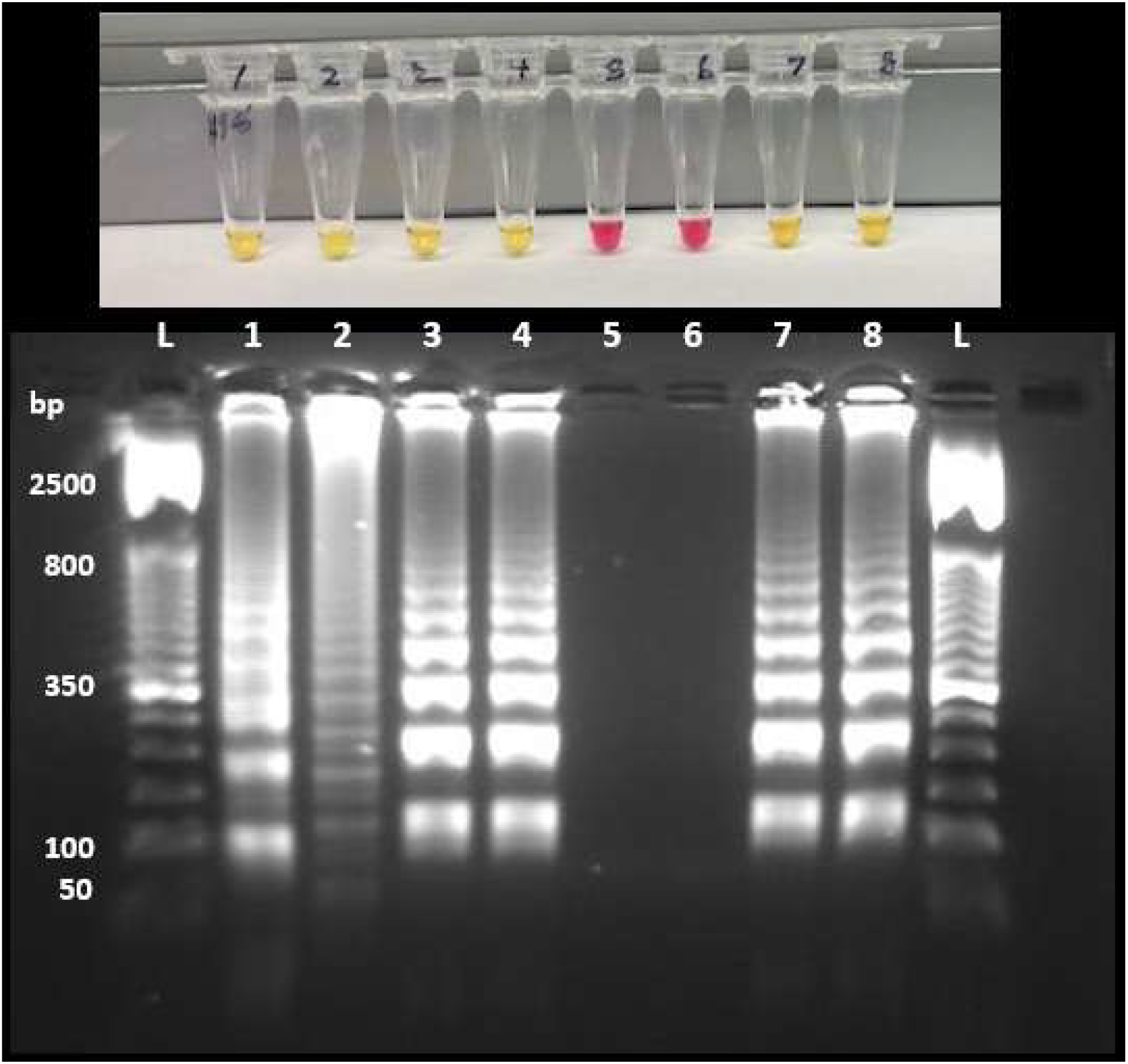
Orthogonal validation of inhibitor-dependent suppression of nonspecific amplification. Representative colorimetric reactions (top) and corresponding agarose gel analysis (bottom) were generated using the M3 primer set in the same polymerase and colorimetric buffer background with or without the nonspecific amplification inhibitor. Without inhibitor (lanes 1–4), NTCs (lanes 1 and 2) changed color and produced amplification products. With inhibitor, NTCs (lanes 5 and 6) remained red and showed no detectable products. Reactions containing heat-inactivated SARS-CoV-2 virions (NR-52286) amplified in both conditions (lanes 3 and 4; lanes 7 and 8), showing that target-dependent amplification was retained.

Without the inhibitor, NTC reactions showed both a color change and detectable amplification products. Inhibitor-containing NTCs remained red and showed no detectable amplification products. Template-containing reactions produced the characteristic LAMP product pattern in both conditions, indicating that target-dependent amplification was retained.

The inhibitor-free NTC products also showed banding patterns distinct from those of the template-derived products, consistent with primer-derived nonspecific amplification. Together, the colorimetric and gel results show that the NTC color change reflected amplification and that the inhibitor suppressed those products while preserving target-dependent amplification. The smart-mug endpoint results were consistent with observations obtained using convection-oven incubation.

The mechanism of inhibition remains to be determined. The experimental results show selective suppression of NTC amplification while target-dependent amplification is retained, but they do not establish the molecular interaction responsible for this selectivity. The inhibitor was identified empirically, and further biochemical studies will be required to define its interaction with the polymerase and primer-derived intermediates.

### 3.3. Performance Across Multiple Primer Sets

To assess performance across primer designs, SARS-CoV-2 RNA (500 copies) and NTCs were evaluated by semi-real-time colorimetric LAMP using four independent primer sets (N008, Lamb, M3, and N27) across replicate experiments (Figure 4). The panel included primer sets with different amplification characteristics, including the amplification-prone M3 set.

**Figure 4.**
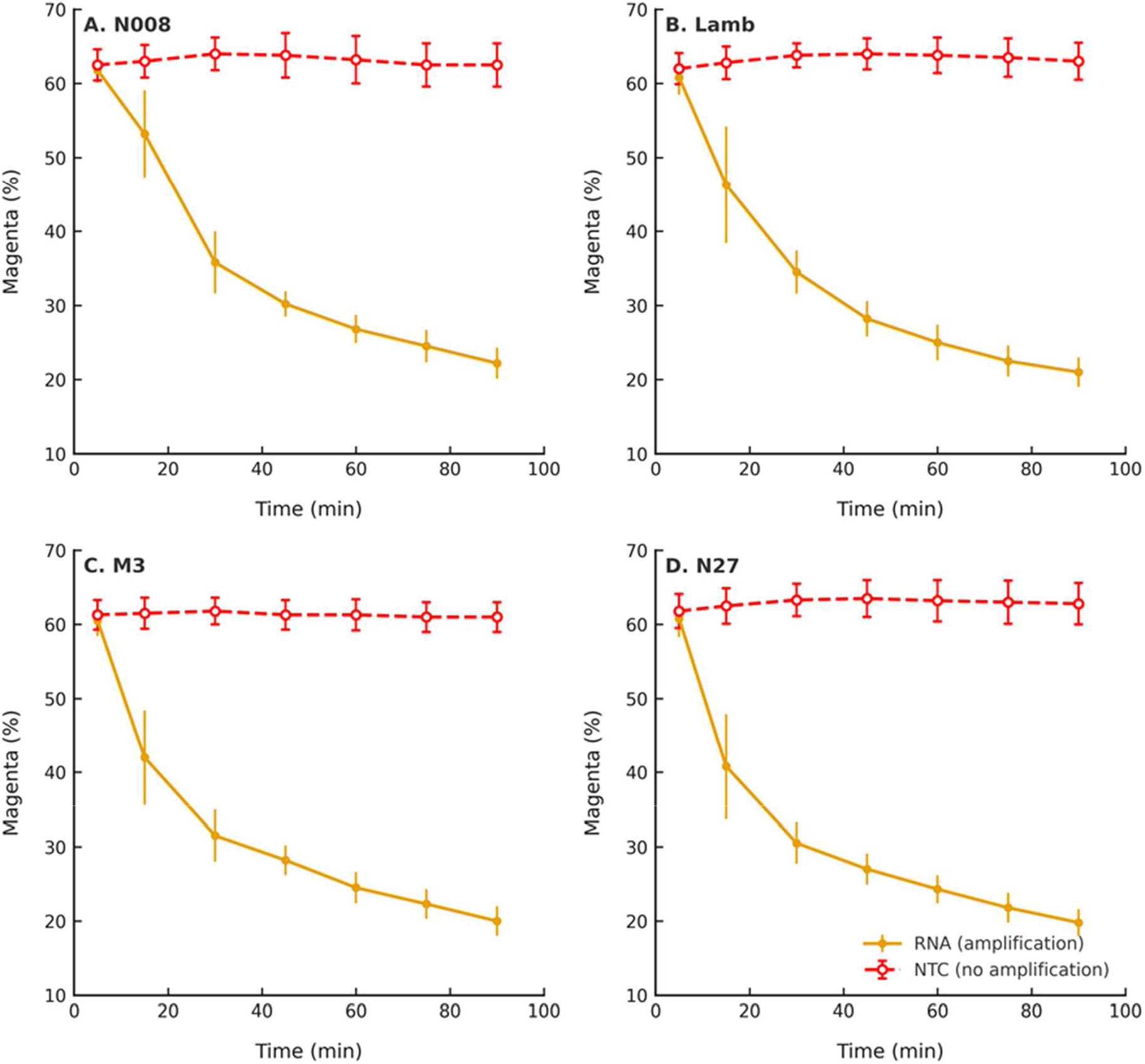
Suppression of primer-driven nonspecific amplification across multiple primer sets. Semi-real-time colorimetric LAMP reactions were performed with four primer sets (N008, Lamb, M3, and N27). Template-containing reactions (solid lines) showed target-dependent color change, whereas TrueLAMP NTCs (dashed lines) remained stable throughout the 90-min incubation. Data are shown as mean ± standard deviation (n=4).

Across all four primer sets, NTC reactions maintained stable magenta intensity throughout the 90-min incubation. Template-containing reactions showed decreasing magenta intensity beginning at approximately 15 min, when a visible color shift became apparent, with reductions exceeding 50% by 30 min.

Template and NTC reactions separated significantly for all four primer sets (Welch’s t-test: p < 0.05 at 15 min and p < 0.0001 after 30 min. Mann–Whitney U test: significant after 30 min). These results show target-dependent amplification with stable NTCs across the four primer designs tested.

### 3.4. Analytical Sensitivity and Specificity

Analytical sensitivity was initially estimated using serial dilutions of SARS-CoV-2 RNA (NR-52347) with the Lamb primer set in a convection oven. Across four independent exploratory experiments, 250 copies per reaction was detected in all four experiments (4/4), whereas 125 copies were detected in only one of four experiments (1/4), and concentrations of 62–16 copies were not detected (Table 1A). Based on these results, 250 copies per reaction was selected as the candidate LOD for formal confirmation.

**Table 1A.** Estimation and confirmation of RT-LAMP limit of detection (LOD) using the Lamb primer set in a convection oven.

|  |  |  |  |  |  |  |  |
| --- | --- | --- | --- | --- | --- | --- | --- |
| RNA (copies/reaction) | 500 | 250 | 125 | 62 | 31 | 16 | 0 |
| Positive Detections | 4/4 | 4/4 | 1/4 | 0/4 | 0/4 | 0/4 | 0/4 |
| LOD Confirmation | RNA (copies) | Positive detections | 95% CI (exact) |  | LOD (copies/reaction) |  |  |
| Oven (65 °C) | 250 | 20/21 (95%) | 77–99% |  | 250 |  |  |

**Table 1B.**
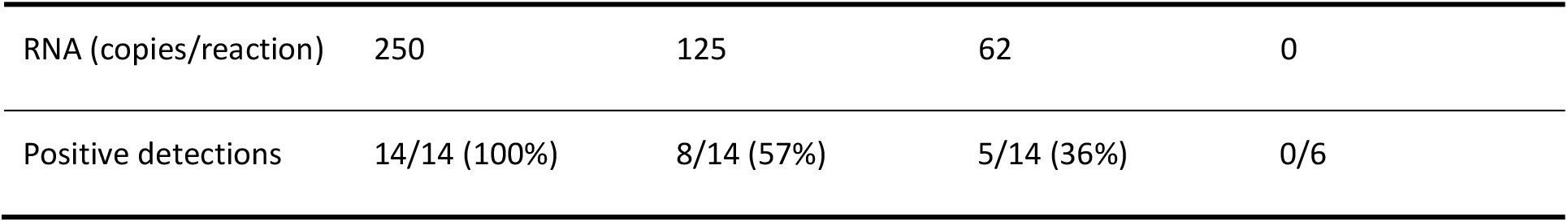
Estimation of RT-LAMP detection sensitivity using the N27 primer set in a convection oven.

|  |  |  |  |  |
| --- | --- | --- | --- | --- |
| RNA (copies/reaction) | 250 | 125 | 62 | 0 |
| Positive detections | 14/14 (100%) | 8/14 (57%) | 5/14 (36%) | 0/6 |

**Table 1C.**
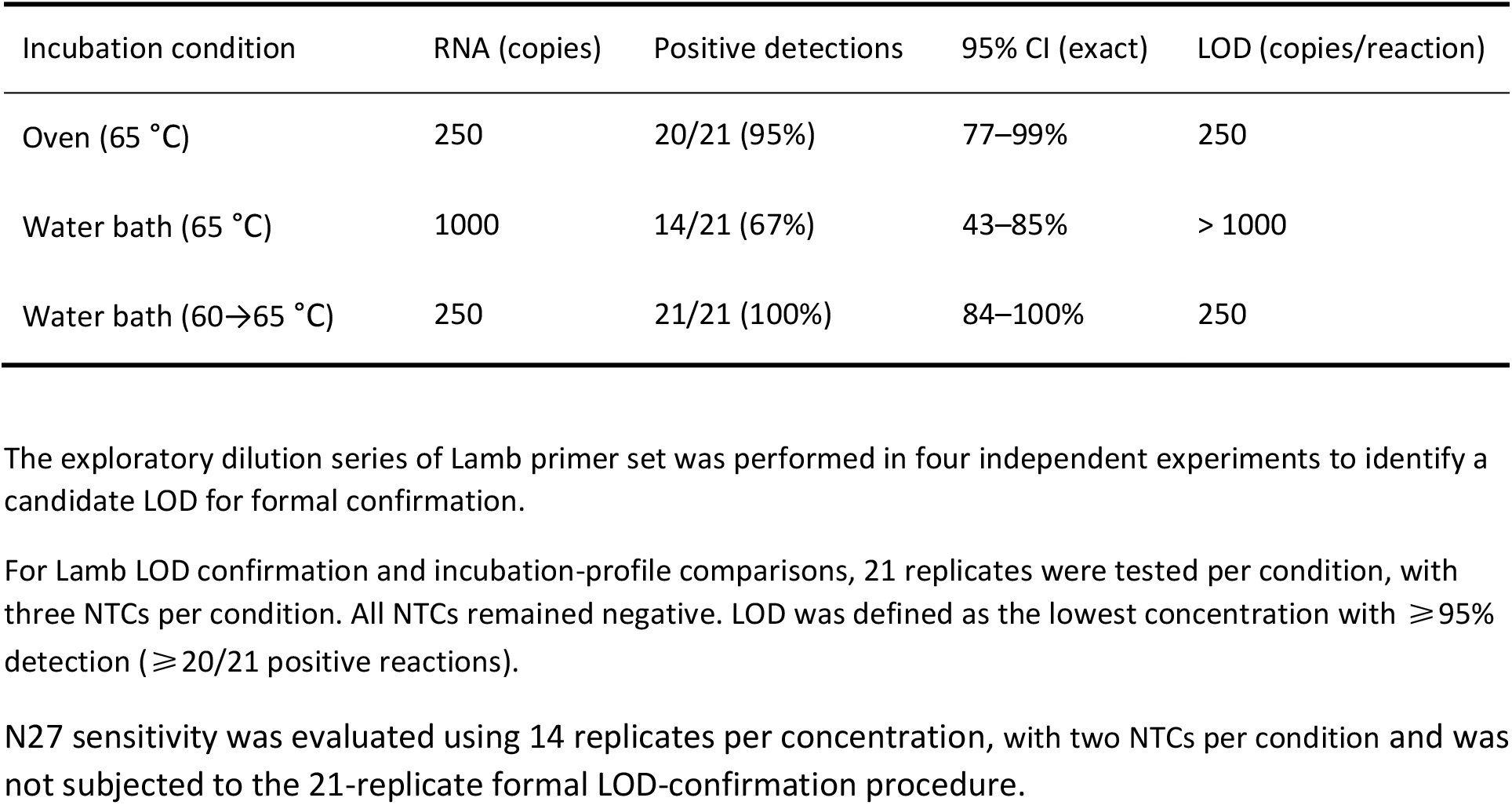
Effect of incubation profile on RT-LAMP limit of detection (LOD)

LOD confirmation using three independent 8-strip experiments yielded 20 positive reactions among 21 tested at 250 copies per reaction (95% detection, 95% CI, 77–99%), while all NTCs remained negative (Figure 5, Table 1A). Based on the prespecified criterion of ≥95% detection, 250 copies per reaction was the LOD for the Lamb primer set under these conditions. Supporting experiments with the N27 primer set showed 14/14 positive reactions at 250 copies per reaction, compared with 8/14 at 125 copies and 5/14 at 62 copies (Table 1B). Although N27 was not subjected to the 21-replicate formal LOD-confirmation procedure used for Lamb, these results independently supported reproducible detection at 250 copies per reaction with a second SARS-CoV-2 primer set. The confirmed Lamb sensitivity is within the range reported for other colorimetric SARS-CoV-2 LAMP assays, for which detection limits vary substantially with primer design, reaction formulation, sample type, incubation time, and LOD criterion [23–25].

**Figure 5.**
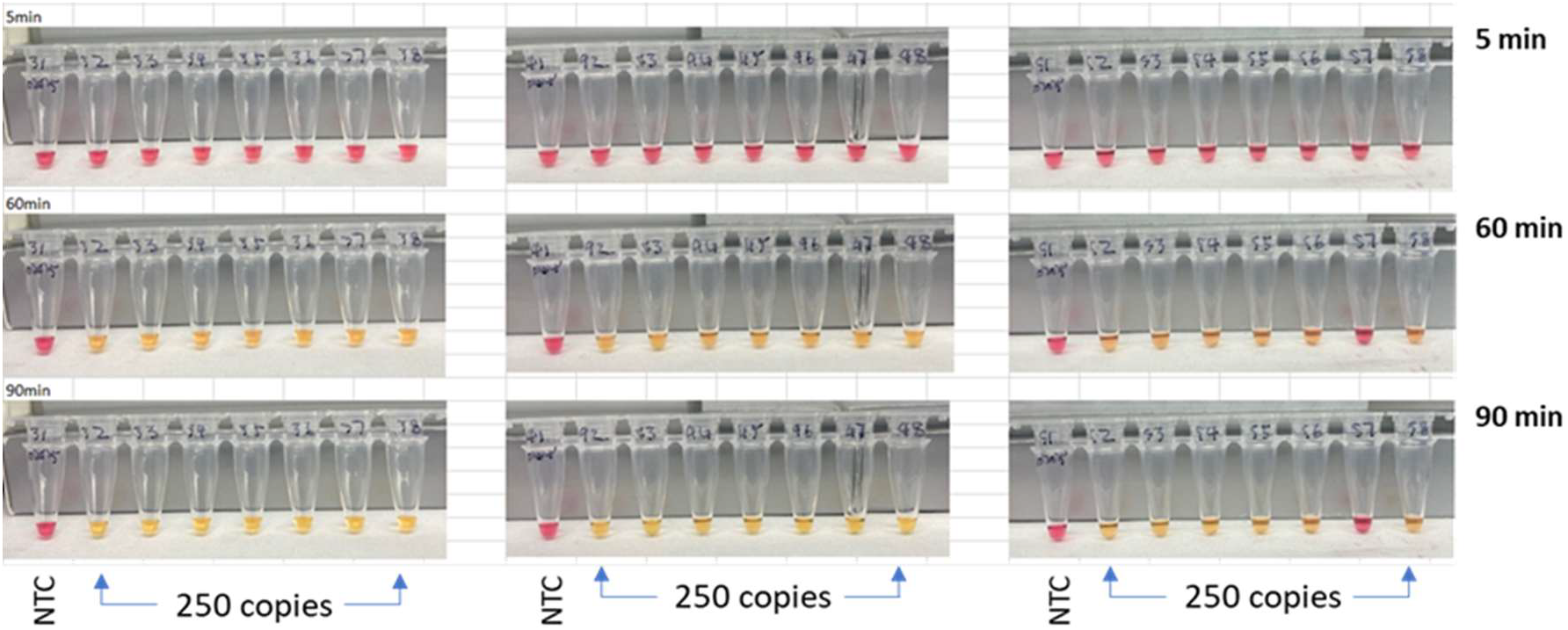
Confirmation of the TrueLAMP limit of detection using the Lamb primer set. Initial dilution experiments showed consistent amplification at ≥250 copies per reaction and inconsistent detection below 250 copies. Three independent 8-strip confirmation experiments yielded 20 positives among 21 reactions at 250 copies (95% detection, 95% CI: 77–99%), establishing 250 copies per reaction as the LOD under the prespecified ≥95% detection criterion.

A total of 65 unique NTC reactions were evaluated: 4 in the inhibitor-comparison experiment (Figure 2), 2 in the gel-validation experiment (Figure 3), 16 across four SARS-CoV-2 primer sets (Figure 4), 3 in the Lamb LOD-confirmation experiment (Figure 5), 24 in the two independent external-laboratory evaluations (Figure 6), 4 in the Lamb exploratory dilution experiments (Table 1A), 6 in the N27 sensitivity experiments (Table 1B), and 6 in the water-bath incubation-profile experiments (Table 1C). All 65 NTC reactions remained negative, corresponding to 100% observed specificity (65/65, exact 95% CI, 94.5– 100%).

**Figure 6.**
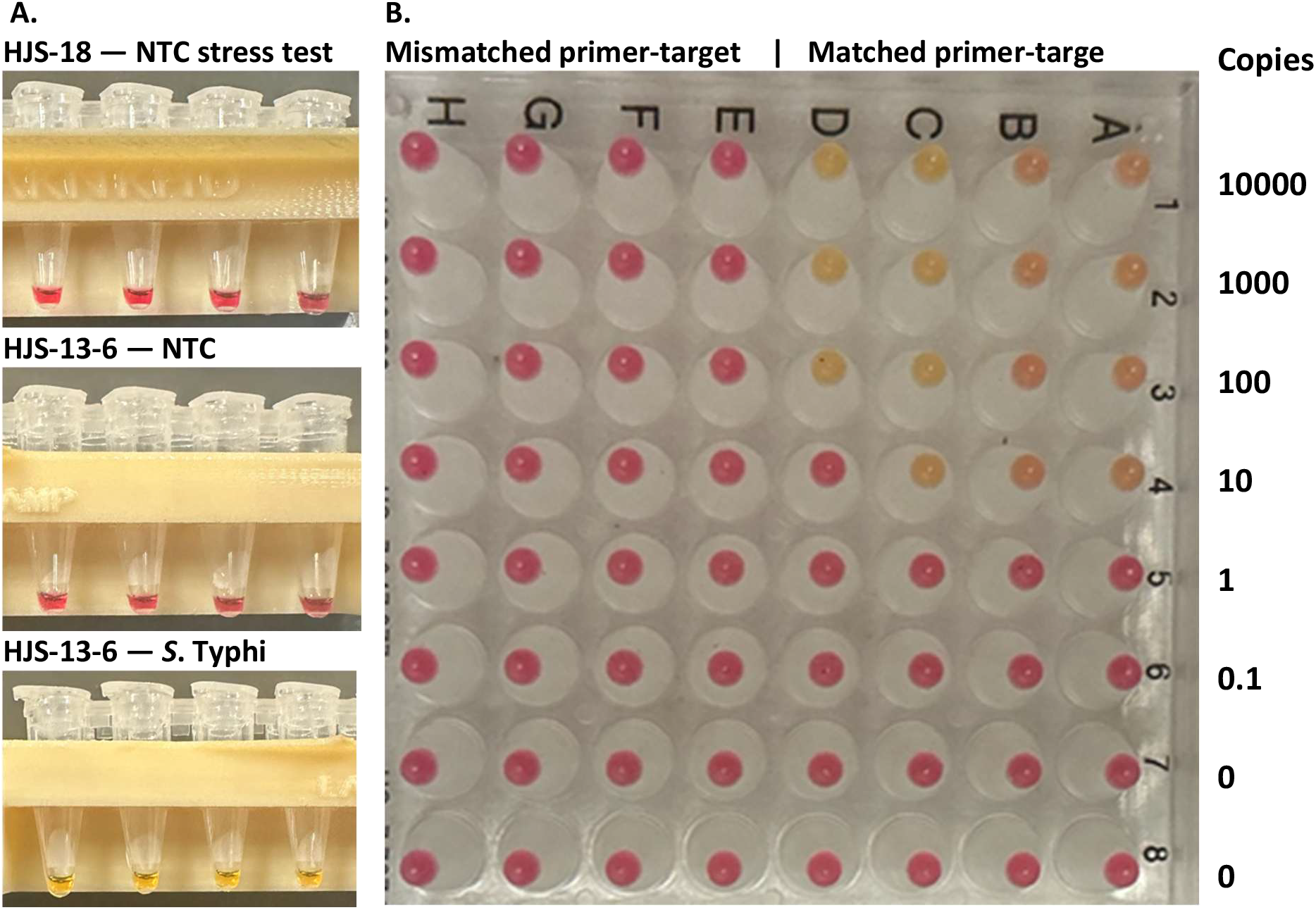
Independent evaluation of TrueLAMP in two external laboratories. (A) The Ellington Laboratory evaluated two previously characterized *S*. Typhi LAMP primer sets. HJS-18, previously reported to show rapid nonspecific NTC amplification, was used specifically as an NTC-suppression challenge. No NTC amplification was observed during the 120-min TrueLAMP incubation. In parallel, HJS-13-6 amplified *S*. Typhi genome (2,000 copies per reaction) with the reactions turning positive/yellow at 30 min while corresponding NTCs remained negative after 120 min. (B) The Mason Laboratory evaluated matched and mismatched primer–template combinations across the indicated template concentrations (0–10,000 copies per reaction). Matched combinations produced positive colorimetric reactions, whereas mismatched combinations remained negative after 120 min.

### 3.5. Two-Step Incubation Ensures Consistent Performance Across Targets, Primer Sets, and Heating Devices

Heating rate affected amplification efficiency. This was most apparent in a 65 °C water bath (Fisher Scientific Model 2LS, Pittsburgh, PA), which equilibrated more rapidly than the convection oven. Under direct 65 °C water-bath incubation, only 14 of 21 reactions (67%) were positive at 1,000 copies, giving an apparent LOD above 1,000 copies, compared with 20 of 21 positives at 250 copies in the convection oven (Table 1C).

Adding an initial 10-min incubation at 60 °C before raising the temperature to 65 °C restored amplification efficiency in the water bath, with 21 of 21 reactions positive at 250 copies (Table 1C). A similar loss of amplification during direct 65 °C incubation, followed by recovery with the two-step profile, was observed with DNA LAMP primer sets that did not require reverse transcriptase. These observations indicate that the initial 60 °C phase improves amplification efficiency for some primer sets before incubation at 65 °C, although the mechanism underlying this effect remains unclear.

Based on these results, we used the two-step profile as the standard TrueLAMP incubation procedure. It provided more consistent amplification across the target types, primer sets, and heating devices evaluated in this study.

### 3.6. Independent Evaluation in External Laboratories

To test whether the specificity findings could be reproduced outside the developing laboratory, TrueLAMP was evaluated independently in the Ellington Laboratory at the University of Texas at Austin and the Mason Laboratory at Weill Cornell Medicine. The two laboratories used additional primer– template systems and complementary experimental designs.

The Ellington Laboratory evaluated the previously characterized HJS-18 and HJS-13-6 primer sets [18]. HJS-18 had previously shown rapid nonspecific amplification in NTC reactions at approximately 30 min and was therefore used specifically as an NTC-suppression challenge. Under TrueLAMP conditions, HJS-18 NTC reactions remained negative throughout the 120-min incubation.

HJS-13-6 retained the rapid target amplification previously reported for this primer set, producing positive reactions with *S*. Typhi complete genomic DNA within 30 min, while the delayed NTC amplification previously observed at approximately 70 min was not detected during 120 min of incubation. Thus, suppression of nonspecific amplification did not require sacrificing rapid target-dependent amplification in this primer system (Figure 6A).

In the Mason Laboratory, matched primer–template combinations produced positive colorimetric reactions across the tested concentration range, whereas mismatched combinations remained negative after 120 min of incubation (Figure 6B). This result independently showed that detectable amplification remained sequence dependent under the conditions tested and that adding nonmatching nucleic acid did not by itself produce a positive reaction.

Together, the external evaluations extend the central findings to different primer sets, target organisms, DNA templates, and experimental settings. Importantly, these studies were performed independently of the developing laboratory and extended the findings from the SARS-CoV-2 RNA/RT-LAMP model to DNA targets. Although broader validation across additional targets and sample matrices will be required, the observed suppression of nonspecific amplification across these systems supports its applicability beyond a single primer set, target, or nucleic-acid template type.

## 4. Conclusion

Direct inhibitor comparisons demonstrated suppression of primer-driven nonspecific amplification, while NTCs remained negative across the primer sets and experimental conditions examined in this study. Target-dependent amplification was retained. Agarose gel electrophoresis confirmed that the color change observed in inhibitor-free NTCs corresponded to primer-driven nonspecific amplification products rather than an indicator artifact. Independent evaluations in two external laboratories extended these observations to additional primer–template systems and DNA targets.

Stable negative reactions provide a wider endpoint-reading window for colorimetric LAMP and reduce dependence on narrowly timed interpretation. TrueLAMP also operated with simple heating and imaging configurations, including a water bath, convection oven, and temperature-controlled smart mug. The present results support suppression of primer-driven nonspecific amplification as a practical way to improve LAMP reliability while retaining a simple isothermal workflow.

## 5. Future perspective

This study primarily used SARS-CoV-2 RNA as a model under controlled laboratory conditions. Independent evaluations in two external laboratories extended the observations to additional primer– template systems and DNA targets. Broader validation is still needed across laboratories, primer designs, target organisms, and clinically relevant specimens to define the generality and reproducibility of TrueLAMP performance.

Future work should also evaluate complete workflows for decentralized testing, including reagent stability, sample preparation, and compatibility with clinically relevant matrices. These studies will help define where TrueLAMP offers practical advantages for point-of-care, public health, veterinary, and environmental nucleic acid testing.

## Author Contributions

Z.C. conceived the study, designed and performed experiments, analyzed the data, and wrote the manuscript.

H.-J.S. and P.C. designed and performed experiments for the independent evaluation of TrueLAMP, analyzed the data, contributed to the corresponding sections of the manuscript, and reviewed and edited the manuscript.

## Acknowledgments

The following reagents were obtained through BEI Resources, NIAID, NIH: SARS-CoV-2 RNA (NR-52347), heat-inactivated SARS-CoV-2 isolate USA-WA1/2020 (NR-52286), and genomic DNA from Salmonella enterica subsp. enterica serovar Typhi strain Ty2 (NR-543). We thank Sanchita Bhadra, Timothy E. Riedel, and Andrew D. Ellington of the Ellington Laboratory, as well as Christopher Mason of the Mason Laboratory, for their support. ChatGPT (OpenAI, San Francisco, CA, USA) was used to assist with language editing and clarity of the manuscript. The authors reviewed and edited all content and take full responsibility for the final text.

## Disclosure statement

Z.C. is the inventor of TrueLAMP and founder of Boltii Diagnostics Inc., which manufactures the TrueLAMP kit. The authors declare no other competing interests.

## Data Availability

All data supporting the findings of this study are included within the article and its Supplementary Materials. The datasets used and/or analyzed during the current study are available from the corresponding author upon reasonable request.

## Funding

This work received no external or institutional funding.

## Notes

### Summary of Updates

This revised version substantially expands the experimental validation and clarifies the scope and interpretation of the study. Major revisions include: Expanded analytical sensitivity studies. The limit of detection (LOD) of the Lamb SARS-CoV-2 primer set was established through four independent exploratory dilution experiments followed by formal confirmation, with 20 of 21 reactions positive at 250 RNA copies per reaction (95% detection; 95% CI, 77-99%). Additional testing with the N27 primer set showed reproducible detection at 250 copies per reaction (14/14 positive), providing supporting sensitivity data with a second primer set. Expanded specificity assessment. Across the experiments reported in the revised manuscript, all 65 unique no-template control (NTC) reactions remained negative, corresponding to 100% observed specificity (65/65; exact 95% CI, 94.5-100%). Independent external validation. TrueLAMP was evaluated independently in two external laboratories using additional primer-template systems. Testing in the Ellington laboratory included previously characterized Salmonella Typhi LAMP primer sets, including a primer set prone to rapid nonspecific amplification. Testing in the Mason laboratory used matched and mismatched DNA primer-template combinations. These experiments extended the findings beyond SARS-CoV-2 RNA/RT-LAMP to DNA targets and independently supported stable NTCs and sequence-dependent amplification. Together, these revisions provide substantially broader experimental support for suppression of primer-driven nonspecific amplification while demonstrating retention of target-dependent amplification across multiple primer sets, target types, and laboratory settings.

