## Supplemental Table 1 for "Suppressing Primer-Driven Nonspecific Amplification in LAMP Using TrueLAMP"

**Supplementary Table S1. Primer sequences used in this study**

| **Set** | **Name** | **Sequence (5′ → 3′)** | **Reference** |
|---|---|---|---|
| Lamb | F3 | TCCAGATGAGGATGAAGAAGA | Lamb 2020 |
| | B3 | AGTCTGAACAACTGGTGTAAG | |
| | FIP | AGAGCAGCAGAAGTGGCACAGGTGATTGTGAAGAAGAAGAG | |
| | BIP | TCAACCTGAAGAAGAGCAAGAACTGATTGTCCTCACTGCC | |
| | LF | CTCATATTGAGTTGATGGCTCA | |
| | LB | ACAAACTGTTGGTCAACAAGAC | |
| N27 | F3 | CCCTCAGATTCAACTGGCAGTA | Huang 2022 |
| | B3 | TTCATTTTACCGTCACCACCAC | |
| | FIP | GAGAGCGGTGAACGGGCGCGATCAAAACAACG | |
| | BIP | TCGAGGACAAGGCTTCGGTAGTAGCCAATTTGGTC | |
| | LF | GACGCAGTATTATTGGGTAAACC | |
| | LB | CCAATTAACACCAATAGCAGTCCA | |
| M3 | F3 | GCTATCGCAATGGCTTGTCTTG | Huang 2022 |
| | B3 | GAAAGCGTTCGTGATGTAGCAA | |
| | FIP | CAGAATAGTGCCATGCCATGTGGTCATTCAATCCAGA | |
| | BIP | AGAAAGTGAACTCGTCGTCCTAGATGGTGTCCAGCAA | |
| | LF | GGCACGTTGAGAAGAATGTTAGT | |
| | LB | GATCCTTCGTGGACATCTTCGT | |
| N008 | F3 | *Proprietary* | Boltii |
| | B3 | *Proprietary* | |
| | FIP | *Proprietary* | |

| Set | Name | Sequence (5′ → 3′) | Reference |
|---|---|---|---|
| | BIP | *Proprietary* | |
| | LF | *Proprietary* | |
| | LB | *Proprietary* | |
| HJS-13-6 | F3 | GAGATGATCACATCATCCATAC | Seo 2026 |
| | B3 | TATTCTCAAGCGCGGGAA | |
| | FIP | GCAACCCTCTGCTTAGATAAAAGTTACACACACTTAGCTTGAGATC | |
| | BIP | TTGGAGATTGAACAAAACCAAAAGGCATTTTCAACTGTAAGAAACTTGTC | |
| | LF | TACTACGCCGATTGAACAAACATCA | |
| | LB | AGGACATGCTTATGTGGACTATGG | |
| HJS-18 | F3 | TCATCCATACACACACACTT | Seo 2026 |
| | B3 | TGATCTAAAGAGCTACTTCACC | |
| | FIP | GCAACCCTCTGCTTAGATAAAAGTTAGCTTGAGATCATGATGTTTGT | |
| | BIP | CAAAAGGGAGGACATGCTTATGTTATTCTCAAGCGCGGGAA | |
| AMV | F3 | CGCGAACCTGAATGAAGCA | Mason Lab |
| | B3 | GCCGATTGGCGAAATCAAC | |
| | FIP | GATGAGGGCAGGATCGCCACCTTCATTTGGTGACCCCGA | |
| | BIP | GACGGCTCTCCAGGCGTTTAGATGGTCCCTGCGTGATGT | |

| **Set** | **Name** | **Sequence (5′ → 3′)** | **Reference** |
|---|---|---|---|
| | LF | GCCGACCACTATTCCCTAACGA | |
| | LB | CGGAACCTGCAGGTCCATG | |
| SHV | F3 | AGTATGGGCAACAACTCA | Mason Lab |
| | B3 | GTGTAATGGCATTGTCTGG | |
| | FIP | CGTCTTATGTAGGACCATTTTGGCTTTCGGTGTTGCTAGAGTC | |
| | BIP | GCACAACCAAGTGTCGATCTTACGGATTTGAGACATATTGGGGAA | |
| | LB | TTGGTGGGGTCAAATTCACAGT | |

**Supplementary Protocol S1. TrueLAMP Protocol**

**Overview**

This protocol describes setup, incubation, and endpoint detection for colorimetric loop-mediated isothermal amplification (LAMP) using the TrueLAMP polymerase and buffer system. The formulation contains an inhibitor designed to suppress primer-driven nonspecific amplification while preserving target-dependent amplification. The protocol is intended for purified nucleic acid templates or sample types shown to be compatible with phenol-red-based colorimetric detection.

**Materials and Reagents**

- TrueLAMP polymerase
- 2x TrueLAMP buffer
- 10x LAMP primer mix (see Supplementary Table S1)
- Template nucleic acid (RNA or DNA)
- Nuclease-free water
- 0.2 mL 8-strip PCR tubes
- Temperature-controlled incubator capable of maintaining 60–65 °C

**Master Mix for one reaction**

Determine the number of reactions required, including positive controls and NTCs. Prepare the master mix with approximately 10% excess volume to compensate for pipetting and dispensing losses.

- 2x buffer: 5.0 µL
- 10x primer mix: 1.0 µL
- Polymerase: 0.05 µL
- Water: 3.0 µL

**Reaction Assembly**

1. Dispense 9 µL master mix into each tube.
2. Add 1 µL template to each test reaction and 1 µL nuclease-free water to each NTC.
3. Vortex to mix thoroughly, then centrifuge briefly to collect the reaction at the bottom of the tube.

**Incubation**

60 °C for 10 min, followed by 65 °C for 30–80 min.

**Detection**

Positive reactions: yellow/orange. Negative reactions and NTCs: red.

For smartphone-based color measurement, sample the region immediately below the liquid meniscus using the smallest available aperture. Record either minimum green intensity in RGB mode or maximum magenta intensity in CMYK mode.

### Practical Considerations and Troubleshooting

#### NTC becomes positive

First verify reaction preparation and incubation conditions. Accurate reagent volumes are important because changes in buffer composition can affect suppression of nonspecific amplification. Confirm the incubation temperature and uniform heating at the tube position. Primer sets should also be evaluated empirically because their propensity for nonspecific amplification varies.

#### Positive reaction does not amplify

Confirm template integrity and concentration and verify that the primer set is functional with the intended target. Establish detection limits empirically for each primer set and sample type. If amplification is inefficient with direct high-temperature incubation, use the recommended two-step profile of 60 °C for 10 min followed by 65 °C.

#### Reaction requires longer than 30 min at 65 °C

Amplification time depends on primer set, target concentration, and sample composition. Many primer sets produce a clear endpoint after approximately 30 min at 65 °C, but slower reactions or low-copy targets may require longer incubation. Continue incubation within the range validated for the assay.

#### Which heating device should be used?

Uniform temperature control is more important than the specific heating device. A temperature-controlled water bath provides rapid and uniform heat transfer. A temperature-controlled smart mug can be used for endpoint testing when the tubes remain adequately immersed and the temperature at the tube position is verified. A convection oven is also suitable and allows semi-real-time imaging through a viewing window.

#### Can unprocessed biological samples be added directly?

Establish compatibility for each sample type. Phenol-red colorimetric detection depends on reaction pH, so samples with substantial buffering capacity, extreme pH, high ionic strength, or other interfering components may alter the colorimetric readout. Purified DNA or RNA is recommended when sample compatibility has not been established.

#### Can TrueLAMP polymerase and 2x buffer be premixed for storage?

Prepare the complete reaction mixture fresh. Storage of TrueLAMP polymerase after combination with the 2x buffer has not been validated for this protocol. Begin amplification promptly after reaction assembly.

#### How long can endpoint results be read after incubation?

Visual amplification results are often apparent after approximately 30 min and can remain stable for many hours under the conditions tested. Interpret endpoints within the validated conditions of the specific assay rather than assuming indefinite stability
